# Resistance Training Reshapes the Gut Microbiome for Better Health

**DOI:** 10.1101/2025.08.13.670057

**Authors:** Daniel Straub, Till Englert, Antonia Beller, Josua Stadelmaier, Mark Stahl, Joachim Kilian, Jens Borzym, Carola Rotermund, Tanja Akbuga-Schön, Sabrina Krakau, Stefan Czemmel, Sabine Weiler, Marc Pettenkofer, Jörg Pettenkofer, Ulli Maser, Sascha Dammeier, Andreas M. Nieß, Markus D Enderle, Sven Nahnsen

## Abstract

**Objectives:** The gut microbiome plays a critical role in metabolism, immunity, and aging. While endurance training has been shown to beneficially modulate the microbiome, the effects of resistance training remain less clear, with some studies reporting minimal changes. This project aims to investigate whether structured resistance training elicits significant changes in gut microbiome composition and diversity in sedentary, healthy adults.

**Methods:** 150 participants completed an 8-week supervised resistance training program. Session-level training data, including weights and repetitions, were recorded alongside metrics like load and compliance. Fecal samples were collected throughout the study period at designated timepoints for 16S rRNA gene amplicon sequencing to assess microbiome composition and for metabolomics analyses to evaluate microbial metabolic activity.

**Results:** No differences in microbial diversity were observed, and there were no significant changes in microbial community composition or fecal metabolomics across all participants post-training. However, within-individual microbial community changes significantly correlated with strength improvement, and significantly stronger shifts in beta diversity were observed in participants with high average strength gains compared to those with smaller gains. In these high responders, differential abundance analysis revealed time-dependent microbial changes, with more taxa enriched or depleted by week 8 of training. Notably, *Faecalibacterium* and *Roseburia hominis*—both associated with a healthier, anti-inflammatory microbiome—were significantly enriched. Many differentially abundant taxa belonged to the *Lachnospiraceae* family.

**Conclusion:** Resistance training drives significant, time-dependent gut microbiome changes, particularly in those demonstrating greater improvements in strength. These shifts mirror endurance training effects and may reflect improved overall health.

**Summary:** While endurance training has been consistently shown to enhance gut microbiome composition, the effects of resistance training remain less well defined, with findings to date being variable. Our results indicate that resistance training can induce meaningful, time-dependent shifts in the gut microbiome, particularly among sedentary individuals who experience substantial strength gains. Notably, we observed enrichment of key health-associated taxa, including *Faecalibacterium* and *Roseburia hominis*, both linked to anti-inflammatory effects and improved gut function. These findings suggest that resistance training may contribute to gut health in conjunction with physical fitness, supporting its broader application in health promotion strategies and future microbiome-focused research.

## INTRODUCTION

The human gut microbiome, comprising trillions of microorganisms, plays a central role in digestion, metabolism, immune function, and overall health [1]. While diet and lifestyle are well-established modulators, growing evidence points to physical activity as a significant influence on gut microbial diversity and composition. Exercise affects key physiological processes such as inflammation and energy metabolism, which are closely linked to the gut microbiome and broader outcomes like physical fitness and aging [2].

The microbiome is increasingly recognized as a marker not only of overall health but also of aging [3,4]. Age-related declines in microbial diversity and beneficial taxa are associated with inflammation, immune dysfunction, and metabolic deterioration [4]. In contrast, a microbiome enriched in bacteria such as *Faecalibacterium, Akkermansia*, and *Roseburia*, known for producing short-chain fatty acids (SCFAs) like butyrate, is linked to improved gut integrity, reduced inflammation, and healthier aging [5]. Endurance training has been consistently shown to promote such a microbial profile, increasing alpha diversity, SCFA production, and gut barrier function [2].

In contrast, the effects of resistance training on the gut microbiome are far less understood. Resistance exercise induces distinct physiological changes, including muscle hypertrophy, increased protein turnover, glycemic control [6] and hormonal adaptations [7]. However, current research offers conflicting findings: while some studies report microbiome shifts correlated with strength performance, others find only minimal or no changes [8,9].

Given the inconsistent findings to date, larger and more detailed studies are needed to determine whether resistance training alone can induce measurable changes in the gut microbiome—particularly changes resembling the health-associated patterns seen with endurance training. To address this gap, we investigated microbiome shifts in a large cohort of previously inactive, healthy adults undergoing a structured resistance training program. Uniquely, we captured detailed session-level data (weights lifted, repetitions performed) to objectively assess training load and adherence. By linking these metrics with longitudinal microbiome and fitness data, this study offers the most comprehensive analysis to date of how resistance training may influence gut microbial diversity and taxa associated with metabolic and immune health.

## MATERIAL AND METHODS

### Training protocol and exclusion criteria

The intervention took place from May 2022 to July 2023 at two “Fitness-und Gesundheitsclub Mapet” locations in Tübingen and Rottenburg, Germany. Sedentary individuals (≥1 year) were screened and enrolled after completing a demographic questionnaire. Participants completed a baseline fitness assessment comprising anthropometric measurements (body weight, body mass index, and body fat percentage assessed via bioelectrical impedance analysis), cardiorespiratory evaluation (VO_2_ max testing), and cardiovascular variables (resting blood pressure and heart rate). Subsequently, participants engaged in an 8-week supervised resistance training intervention (2–3 sessions per week). Follow-up fitness assessments were conducted at weeks 4 and 8. Participants completed pre-test questionnaires covering dietary and fluid intake, medication use and other relevant factors, and were instructed to maintain their usual dietary and fluid intake patterns throughout the study period.

Training was conducted on seven EGYM Smart Strength machines (Seated row, Lat Pull, Chest press, Back trainer, Abdominal trainer, Leg curl, and Leg press) using one of two resistance training programs: a resistance training program aimed at improving general fitness, or muscle-building (Table 1). The equipment enabled standardized training through automated control of range of motion, movement speed, and digitally programmed resistance profiles, including eccentric overload phases and adaptive load adjustments based on user performance. These algorithm-driven protocols provided progressive, individualized stimuli not feasible with conventional gym equipment. Training machines not only guided each session in real time but also automatically recorded detailed exercise data (machine used, program, weight, repetitions, max strength, date) for every workout.

**Table 1:**
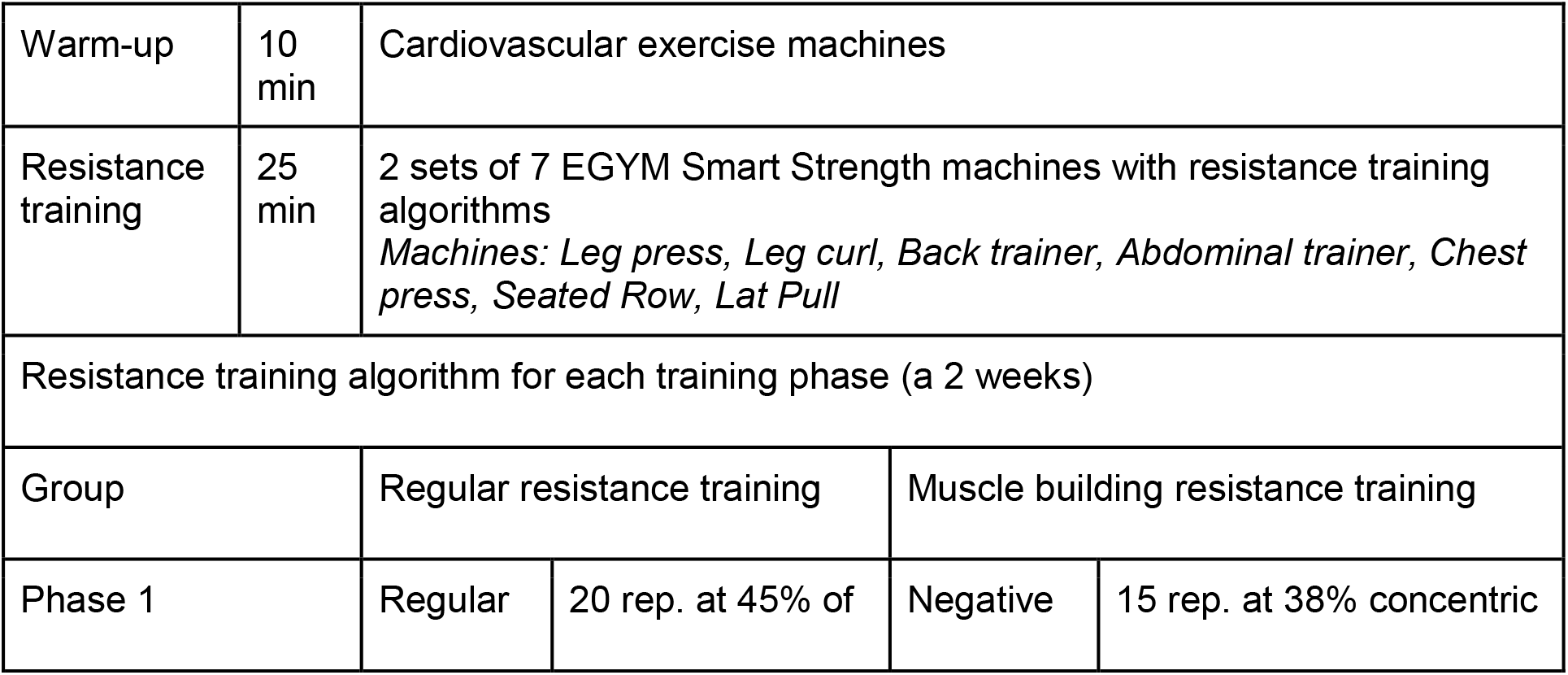
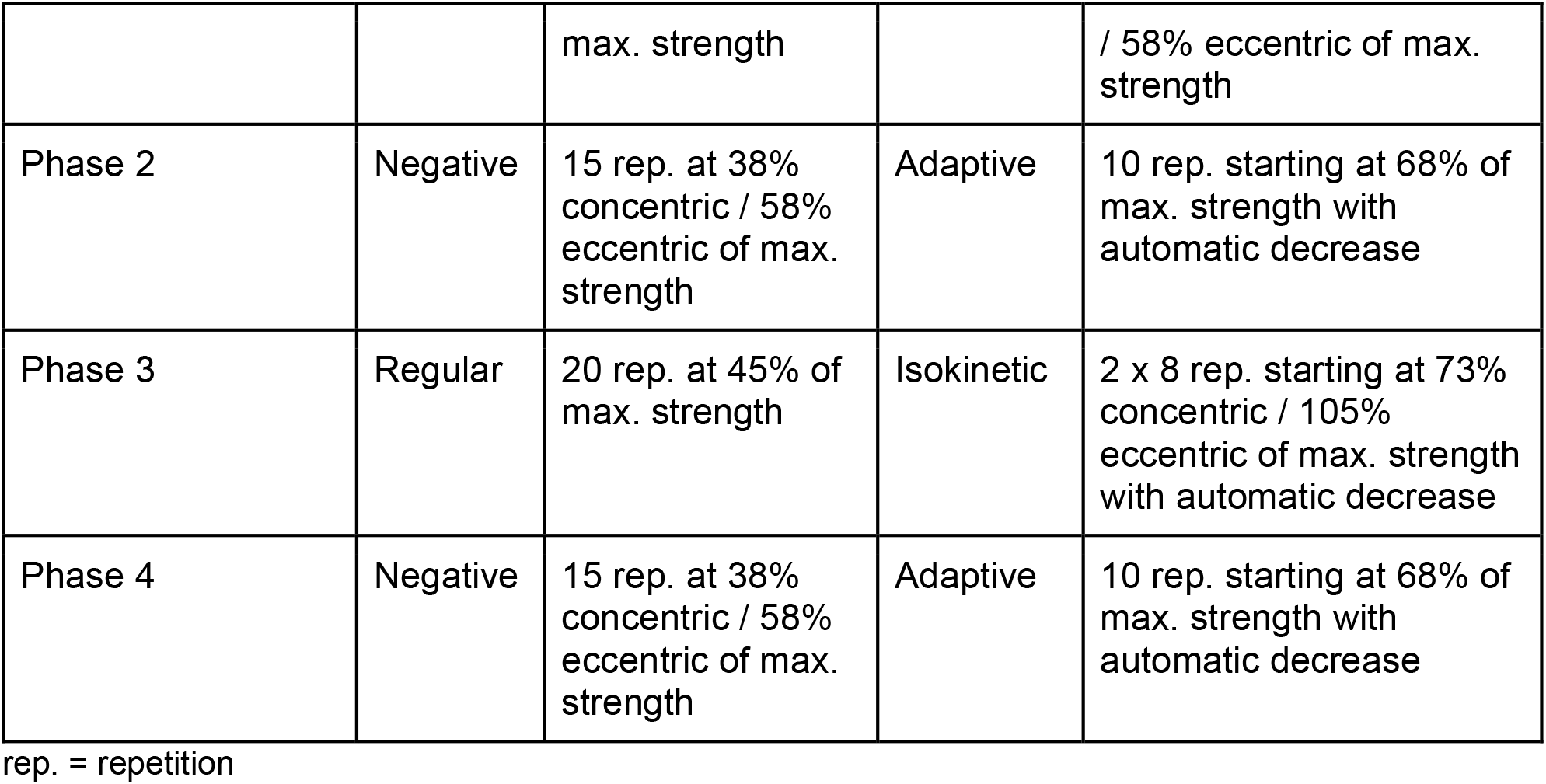
Training protocol with EGYM resistance exercise machines and training algorithms.

|  |  |  |  |  |  |
| --- | --- | --- | --- | --- | --- |
| Warm-up | 10 min | Cardiovascular exercise machines |  |  |  |
| Resistance training | 25 min | 2 sets of 7 EGYM Smart Strength machines with resistance training algorithms<br>*Machines: Leg press, Leg curl, Back trainer, Abdominal trainer, Chest press, Seated Row, Lat Pull* |  |  |  |
| Resistance training algorithm for each training phase (a 2 weeks) |  |  |  |  |  |
| Group |  | Regular resistance training |  | Muscle building resistance training |  |
| Phase 1 |  | Regular | 20 rep. at 45% of | Negative | 15 rep. at 38% concentric |

|  |  |  |  |  |
| --- | --- | --- | --- | --- |
|  |  | max. strength |  | / 58% eccentric of max. strength |
| Phase 2 | Negative | 15 rep. at 38% concentric / 58% eccentric of max. strength | Adaptive | 10 rep. starting at 68% of max. strength with automatic decrease |
| Phase 3 | Regular | 20 rep. at 45% of max. strength | Isokinetic | 2 x 8 rep. starting at 73% concentric / 105% eccentric of max. strength with automatic decrease |
| Phase 4 | Negative | 15 rep. at 38% concentric / 58% eccentric of max. strength | Adaptive | 10 rep. starting at 68% of max. strength with automatic decrease |

rep. = repetition

Of the 205 enrolled participants, 43 did not complete the full 8-week training protocol and were excluded from analysis. Six additional participants were excluded from the analysis due to data quality concerns (antibiotic use or improper stool sample storage). Another six participants were excluded as their samples were used for internal validation of sequencing and metabolomics workflows.

### Strength and Training Metrics

Average strength gains were assessed by calculating the average log_2_ fold change in maximum strength across all exercise machines between two time points.

BioAge Strength is an EGYM metric that estimates the biological age of muscular fitness based on a non-linear model of the typical relationship between age and strength. It uses a large dataset from approximately one million probands including chronological age, sex, and strength relative to body weight to determine whether an individual’s strength is above (younger BioAge) or below average (older BioAge) for their chronological age. The final score is calculated as the average across all exercise machines and can be improved by increasing strength or reducing body weight—but not below the minimum of 21 years.

Activity Points, another EGYM metric, were used to estimate the metabolic equivalent (MET) expended per session. This measure integrates training intensity, excess post-exercise oxygen consumption (EPOC), the mechanical work of each set, and an individualized estimate of the participant’s resting metabolic rate, providing a comprehensive representation of training load.

Training compliance was quantified as the percentage of targeted repetitions actually performed across all sessions, offering an objective measure of adherence to the training protocol.

### Stool sample processing

Stool samples were self-collected by participants using two OMNIgene®-GUT kits (OM-200 for 16S rRNA gene amplicon sequencing and ME-200 for targeted metabolomics) and stored at 4°C before transport at ambient temperature and long-term storage at −80°C.

DNA was extracted using the DNeasy 96 PowerSoil Pro Kit and 16S rRNA V4 regions were amplified using primers 515F/806R. Sequencing was performed on the Illumina MiSeq platform (2 × 250 bp, 69k reads/sample on average). Data processing employed the nf-core/ampliseq pipeline (v2.7.0) [10] as detailed in the supplement.

For metabolomics, samples were prepared, analyzed and quantified via internal standards and dry weight normalization via LC-MS and GC-MS as detailed in the supplement.

### Statistical Analysis

Non-parametric tests (Friedman, Conover post hoc) were used to assess fitness changes over time. Longitudinal trends between training groups were analyzed with *nparLD* in R. Subsets of lowest 20% responders were compared to the highest responders (n=30 each) - based on either maximum strength gain in the leg press, or average strength gain for all 7 exercises, or average decrease of BioAge strength - either using Kruskal-Wallis rank sum test followed by pairwise comparisons using Wilcoxon rank sum test with continuity correction or Kolmogorov-Smirnov tests for distributions. Multiple testing correction was done with the Benjamini-Hochberg method. Significance threshold was set at p<0.05 if not otherwise indicated.

Differential microbial abundance was assessed with ANCOM-BC2 [11] using Dunnett’s test [12] with Holm’s family-wise error rate correction [13] based on phyloseq [14] objects from nf-core/ampliseq, modeling time point as a categorical variable. Extended information is available in the supplement.

### Equity, diversity, and inclusion statement

Our study includes similar numbers of sedetary women and men in the vincinity of Tübingen, Germany. We did not purposefully recruit people from marginalized communities. Our author team included five women and fourteen men, at the time all living in Germany. Author disciplines included bioinformatics, biology, sport management, sports medicine, internal medicine, and we included three junior scholars. We did not examine the effects of race/ethnicity or socioeconomic status. We discussed gender specific limitations on our findings in the discussion.

### Patient involvement

Participants and the public were not and will not be involved.

## RESULTS

### Resistance training rapidly enhances muscle strength

Of the 205 participants initially enrolled, 150 were included in the final analysis (Figure 1A). Table 2 summarizes the demographic and baseline characteristics of the final study cohort.

**Table 2:** Demographic data of participants. Values as percent or mean ± SD.

| Training group | All participants<br>n = 150 | Regular resistance<br>training<br>n = 73 | Muscle-building<br>resistance training<br>n = 77 | p-value |
| --- | --- | --- | --- | --- |
| Gender (male) | 42.6% | 43.7% | 41.6% | 0.927<sup>1</sup> |
| Age (years) | 41.7 ± 11.6 | 43.1 ± 11.4 | 40.5 ± 11.7 | 0.144<sup>2</sup> |
| BMI (kg/m<sup>2</sup>) | 24.5 ± 3.5 | 24.7 ± 3.7 | 24.4 ± 3.4 | 0.683<sup>2</sup> |
| Non-smoker | 76.7% | 78.1% | 75.3% | 0.827<sup>1</sup> |
| Any allergy | 32.0% | 27.4% | 36.4% | 0.317<sup>1</sup> |
| No special diet | 82.7% | 89.2% | 77.0% | 0.094<sup>1</sup> |
| Born in<br>Germany | 82.4% | 87.3% | 77.9% | 0.199<sup>1</sup> |

<sup>1</sup>: Pearson's Chi-squared test with Yates' continuity correction
<sup>2</sup>: Kruskal-Wallis rank sum test

**Figure 1:**
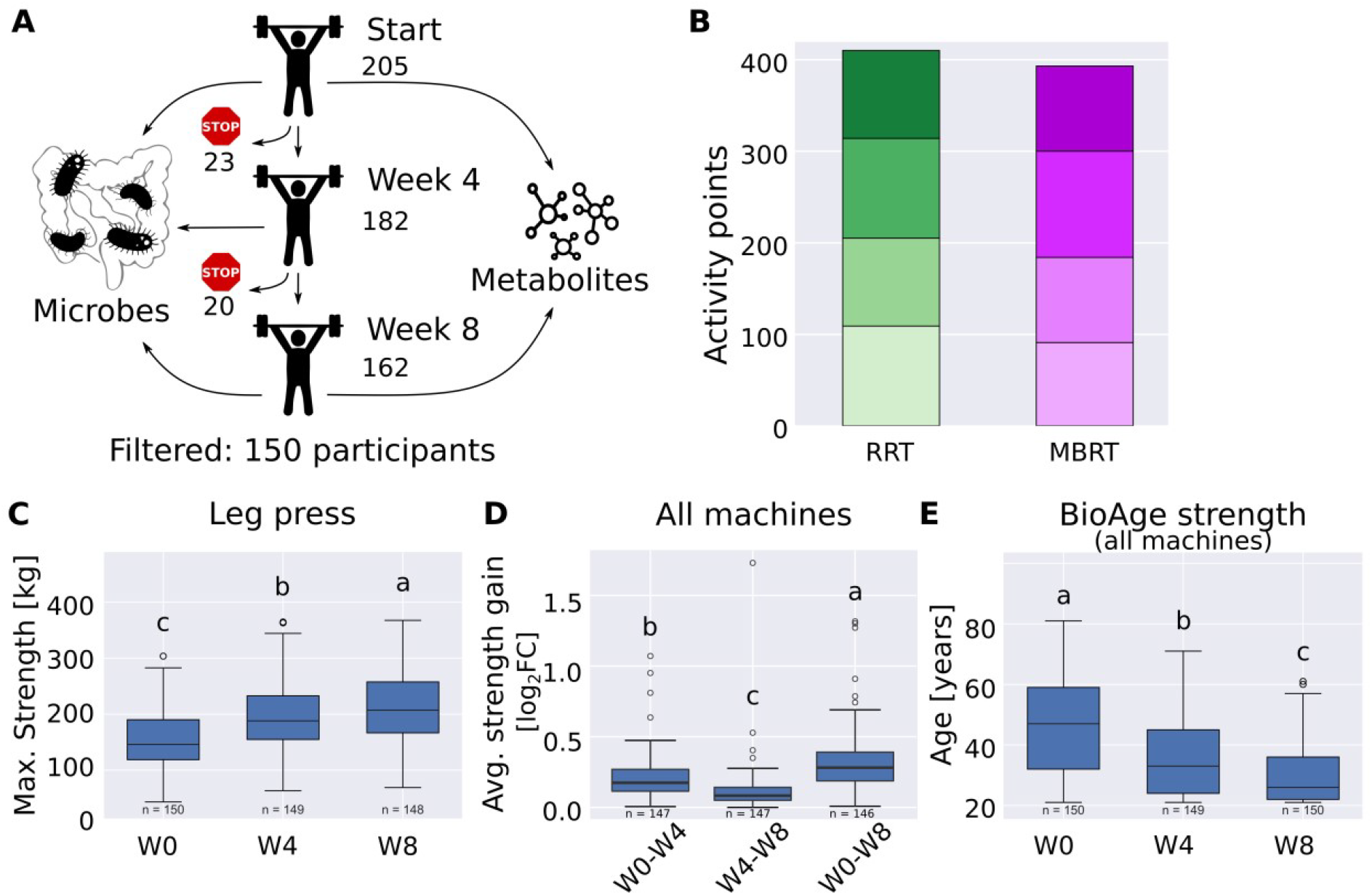
(A) Study overview with number of participants enrolled in the study, with three fitness tests, three gut microbiome samples and two gut metabolome samples per participant. (B) Comparison of activity points (metabolic equivalents minutes) of regular resistance training (RRT) and muscle-building resistance training (MBRT). (C) Maximum strength of leg press over time. (D) Average strength gain over seven exercises. (E) Calculated BioAge Strength over time. log_2_FC: log_2_ fold change, W0: week 0, W4: week 4, W8: week 8. Letters denote p<0.05.

Participants were randomly assigned to one of two training regimens: regular resistance training (higher repetitions, lower weights) or muscle-building resistance training (higher weights, fewer repetitions) (Table 2). The study initially aimed to compare fitness and microbiome responses between the two approaches. Despite differences in training style, both programs were similar in total weight moved and activity points (metabolic equivalent minutes) (Figure 1B). Further, no significant differences in fitness improvements were observed between the groups. As a result, participants were pooled for all subsequent analyses to maximize statistical power.

Strength gains were evaluated using three metrics: (1) maximum strength leg press (kg), (2) average strength gain (average log_2_ fold change across all machines), and (3) BioAge Strength (years). All measures showed substantial improvement within the first four weeks, followed by a smaller increase in the second half of the program. On average, leg press strength rose from 154 ± 54 kg to 217 ± 65 kg (Figure 1C), average strength increased by 24 ± 16% (Figure 1D), and BioAge Strength decreased from 45.8 ± 14.7 to 29.6 ± 9.6 years (Figure 1E).

Beyond strength, participants also experienced statistically significant, though small, reductions in diastolic blood pressure (from 108 ± 17 to 105 ± 17 mmHg) and body fat percentage (from 29.72 ± 8.01% to 29.34 ± 8.30%). However, BMI, resting heart rate, and systolic blood pressure remained unchanged (Suppl. Fig. 1).

### Changes in the Microbial Community is associated with Strength Gain

Resistance training did not significantly alter gut microbial alpha diversity, neither across all participants (Figure 2A) nor within groups stratified by strength metrics. Similarly, no notable changes were observed in beta diversity across these groups, and stool metabolomic profiles remained unchanged (Suppl. Fig. 2). Longitudinal within-subject analyses demonstrated that shifts in microbial community composition were modestly, yet significantly, associated with improvements in strength performance—most notably with average strength gains (Figure 2B).

**Figure 2:**
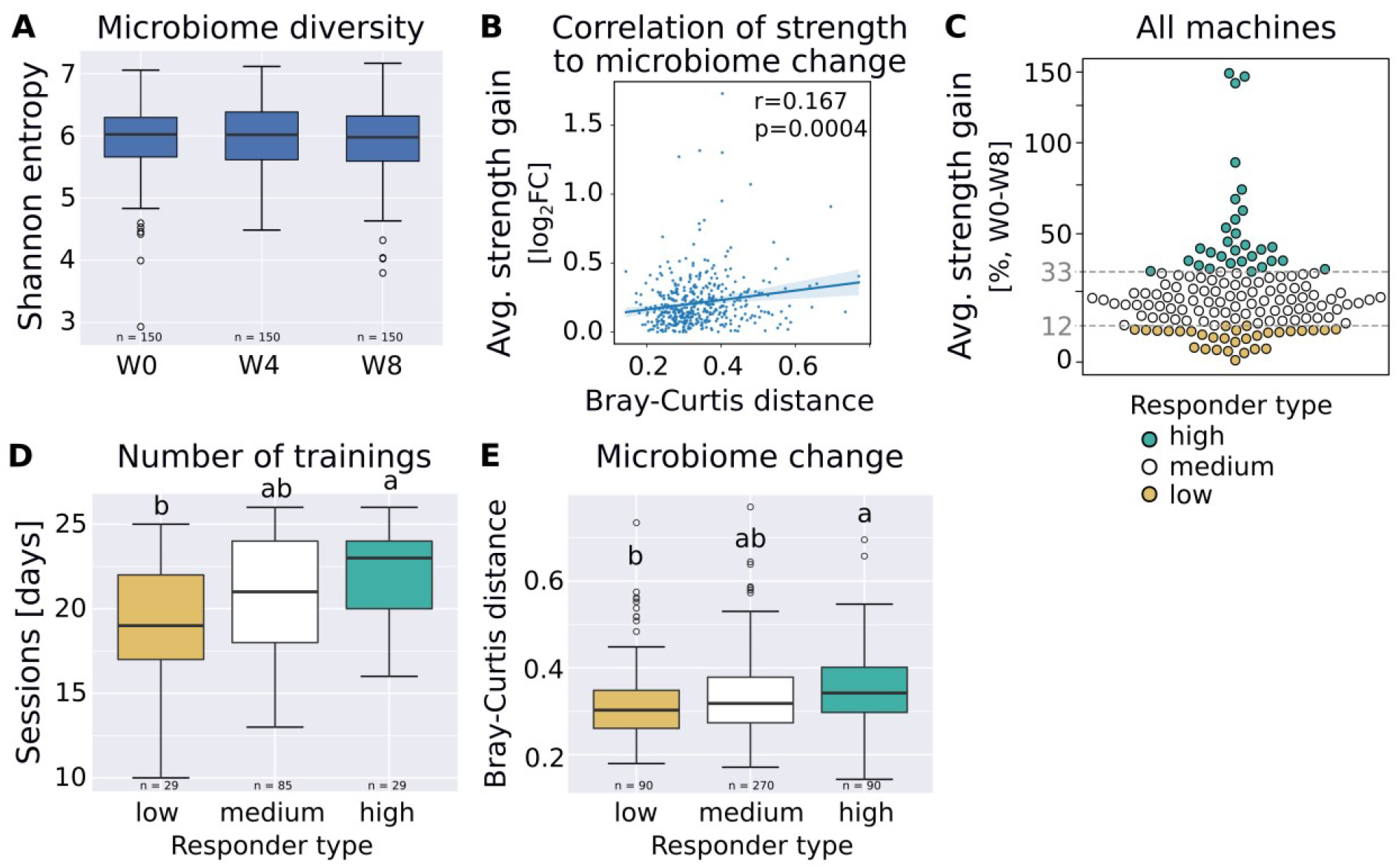
(A) Microbial diversity over the course of the training (p>0.1 for Friedman test followed by pairwise Friedman-Conover test). (B) Pearson correlation between changes of the microbial community and changes in fitness (average strength gain) over 8 weeks. (C) Subsetting participants in low, medium and high responders based on change in average strength: Low responders had less than 12.2% strength gain, high responders more than 33%. (D) Number of training days per participant stratified by responder type (average strength). (E) Microbial community changes (Bray-Curtis distance) within participants stratified by responder type (average strength). log_2_FC: log_2_ fold change, W0: week 0, W4: week 4, W8: week 8. Letters denote p<0.05.

Based on this correlation, participants were stratified into high-responder (HR, top 20%) and low-responders (LR, bottom 20%) across three strength metrics (Fig. 2C), yielding three partially overlapping subsets (Suppl. Fig. 3). Among these, the average strength gain subset showed the least demographic bias, while the BioAge Strength subset showed the most (Suppl. Fig. 4). Dietary changes did not differ significantly between HR and LR groups in any subset (Suppl. Fig. 5).

**Figure 3:**
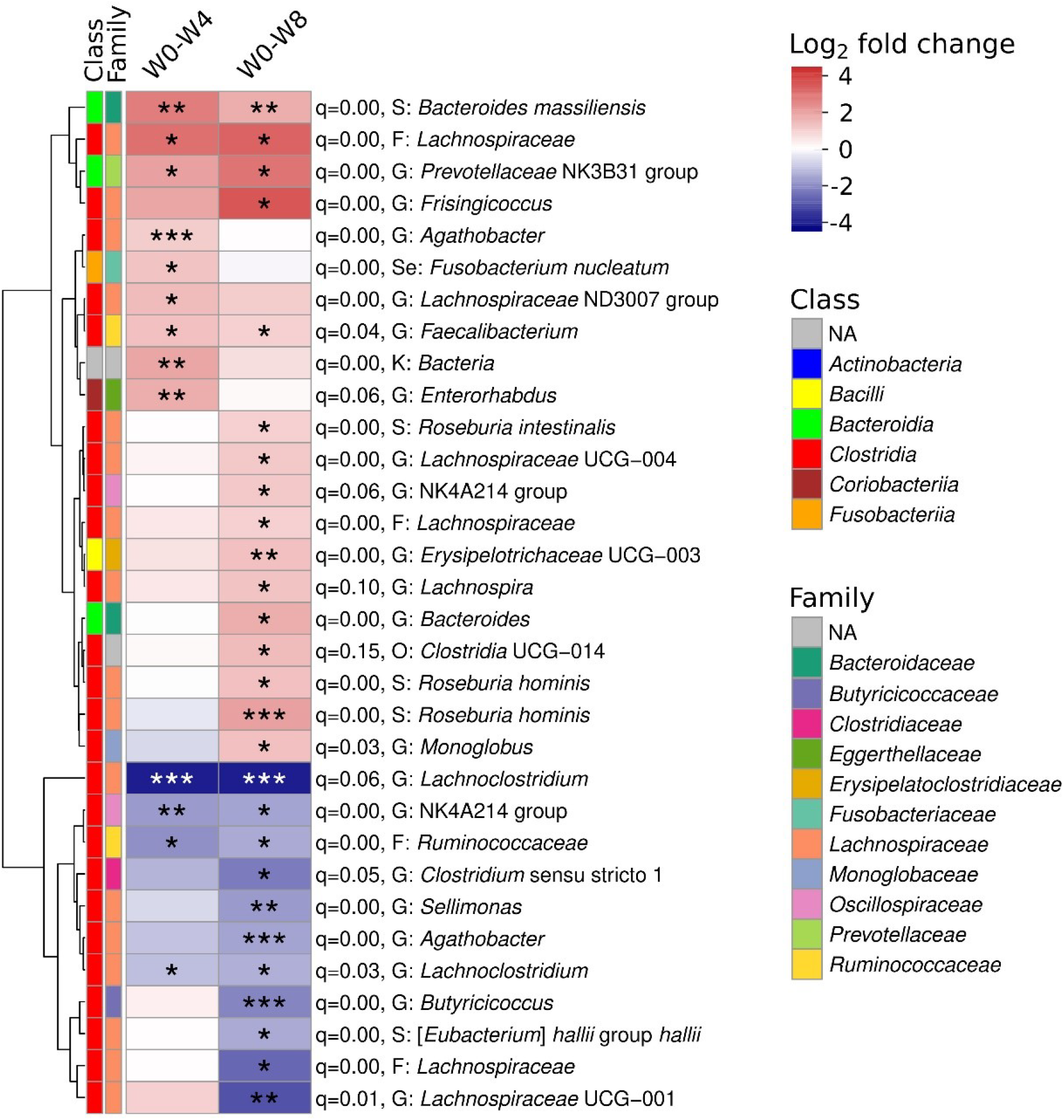
Heatmap of significantly different Amplicon Sequence Variants (ASVs) with taxonomies within high responders (average strength gain) over time. The q-value indicates Benjamini-Hochberg corrected p-values that ASVs were found by chance (permutation test with 100 random subsets of non-high responder participants). For each ASV, class and family are indicated as colours, and the lowest reported informative taxonomic level is indicated as K: Kingdom, O: order, F: family, G: Genus, S: Species, Se: Species with exact sequence match. W0: week 0, W4: week 4, W8: week 8. ANCOM-BC2 (formula: asv ∼ timepoint + (1|participant), dunnet test) with p≤0.05 and log_2_ fold change ≤-1 or log_2_ fold change ≥1. *: p≤0.05, **: p≤0.001, ***: p≤0.00001.

The number of training days differed significantly between HR and LR only in the average strength gain subset (20.7 vs. 19.1 sessions; Figure 2D), while training compliance was similar across all groups. A Random Forest model using baseline fitness- and metadata identified strength measures as the strongest predictors of training response; microbial and metabolomic data were not predictive (Suppl. Fig. 6).

Finally, within-individual beta diversity distances differed significantly between HR and LR in the average strength gain subset (Figure 2E), but not in the other two strength-based groupings.

### Differential Microbiome Changes in High-Responder

As the correlation of longitudinal within-subject shifts in microbial community composition with strength performance was most pronounced in the average strength gain group, this subset was selected for further differential abundance analysis (DAA), focusing on HR. To ensure robustness, a permutation-based approach was used: random subsets of non-high responders (not classified as high responders by any metric) were also subjected to DAA and microbial changes in HR with a q-value < 0.05 were considered unlikely to be due to chance.

DAA revealed distinct microbial shifts in HRs following resistance training. After four weeks, nine Amplicon Sequence Variants (ASVs) were significantly enriched and four were depleted compared to control subsets. By week eight, these differences became more pronounced, with sixteen ASV increased and eleven decreased in abundance. Notably, a large proportion of differentially abundant sequences belonged to the *Lachnospiraceae* family. However, no consistent phylogenetic pattern emerged, as both increases and decreases occurred across diverse microbial lineages.

Several ASVs were consistently and significantly enriched at both time points, including ASVs that were associated to *Faecalibacterium, Bacteroides massiliensis, Lachnospiraceae*, and the *Prevotellaceae* family. Conversely, consistently and significantly depleted ASVs included *Lachnoclostridium*, along with sequences from the *Oscillospiraceae* (NK4A214 group) and *Ruminococcaceae* families. Notably, an ASV classified as *Roseburia hominis* showed the most significant increase, while *Agathobacter* and *Butyricicoccus* were the most significantly depleted, changes that reached significance only at week eight (Figure 3).

## DISCUSSION

This study provides evidence that structured resistance training can induce significant and reproducible shifts in gut microbiome composition, particularly in individuals exhibiting pronounced adaptations in muscle strength. By utilizing a large cohort of 150 previously inactive but otherwise healthy adults and integrating digitally controlled resistance training protocols, individual strength trajectories were objectively linked to microbial dynamics. A key methodological strength was the use of digitally controlled resistance training equipment, which enabled individualized and progressively adjusted training stimuli. These standardized protocols likely enhanced the precision and consistency of training across participants, potentially contributing to the observed association between strength development across all exercises and microbial changes. Participants with lower baseline strength demonstrated the greatest relative improvements, consistent with the principle of diminishing returns in resistance training [15,16]. These findings support the potential of resistance training, particularly when delivered through adaptive modalities, as a modulator of host–microbe interactions.

The absence of significant changes in alpha diversity following resistance training contrasts with findings from endurance-based exercise interventions, where increases in microbial richness and evenness are commonly reported [2,17–19]. This suggests that alpha diversity may be less responsive to resistance training stimuli [9]. However, shifts in beta diversity among participants with greater strength gains indicate that compositional remodeling does occur and may depend on surpassing a physiological threshold, consistent with a dose– response relationship [20].

The enrichment of taxa such as *Faecalibacterium* and *Roseburia hominis*, both producers of SCFAs with anti-inflammatory properties, suggests that resistance training may foster a gut microbial profile conducive to metabolic health and immune regulation [5]. SCFAs have been implicated in maintaining gut barrier integrity, regulating glucose homeostasis [6], and exerting systemic anti-inflammatory effects [2]. These findings are consistent with those reported by Cullen et al. [8], who observed increased abundance of *Roseburia* and a non-significant trend toward higher levels of *Faecalibacterium prausnitzii* following a 6-week resistance training program in young adults with overweight and obesity. Their results further support the potential of resistance training to promote SCFA-producing taxa, with possible implications for improved insulin sensitivity [6] and cardiometabolic health [8]. Notably, similar microbial adaptations, including increases in SCFA-producing genera such as *Roseburia*, have been documented in response to endurance training, highlighting that despite differing exercise modalities, both resistance and endurance training may converge on shared microbiome-mediated pathways that enhance host metabolic and immune function [2,21].

The modulation of *Lachnospiraceae*, a diverse bacterial family with both beneficial and potentially pathogenic members, illustrates their complex, context-dependent role in microbiome adaptation to physical activity. While some murine studies link certain strains to reduced endurance, others report performance-enhancing effects from taxa like *Coprococcus eutactus* [e.g. 22–24]. Importantly, all observed shifts in microbial composition—including the enrichment of *Faecalibacterium* and *Roseburia hominis*, as well as changes within the *Lachnospiraceae* family—occurred in the absence of dietary modification, indicating that resistance training alone may act as an independent driver of gut ecosystem remodeling.

Several mechanisms have been proposed to explain how these microbial shifts may influence muscular adaptations to resistance training. SCFAs—particularly butyrate, produced via bacterial fermentation of dietary fiber—have demonstrated beneficial effects on muscle metabolism and function [25]. The gut microbiota also play a key role in amino acid biosynthesis and metabolism, which are essential for muscle protein synthesis [26,27], as well as in regulating energy homeostasis, which can impact muscle anabolism and recovery [28]. These interconnected pathways suggest that the gut microbiome may contribute to the physiological adaptations induced by resistance training, although further mechanistic studies are warranted to clarify these relationships.

Despite these compositional changes in the microbiome, no significant alterations were detected in the stool metabolomic profile, an observation that contrasts with previous reports demonstrating a correlation between gut microbiome composition and metabolomic output [29]. This discrepancy may reflect subtle or localized functional changes, limitations of the targeted metabolomics approach, or functional redundancy within the microbiome. It is also possible that transient metabolic responses were missed due to the exclusive use of long-term sampling; short-term assessments may reveal dynamic shifts not captured in this study.

It is also important to consider that stool metabolomics predominantly reflect luminal microbial activity and may not fully capture host–microbe metabolic interactions occurring at the mucosal interface or systemically. Resistance training may influence host metabolism through pathways not reflected in fecal metabolites, including muscle-derived signaling molecules, systemic inflammation, or shifts in energy substrate utilization. Taken together, these findings emphasize the need for future investigations to integrate multi-omics approaches, such as plasma metabolomics and metatranscriptomics, to more comprehensively characterize the functional impact of exercise-induced microbiome alterations.

From a translational perspective, these findings carry important implications for public health and clinical practice. Resistance training is already recommended for musculoskeletal health and metabolic disease prevention; our results suggest it may also serve as a non-pharmacological strategy to enhance gut health. This opens avenues for integrating resistance training into holistic health promotion programs, particularly for populations at risk of chronic inflammation, metabolic syndrome, or age-related decline in gut function.

Despite its strengths, this study has limitations. The lack of a non-exercising control group precludes definitive causal inference, and the generalizability of findings to clinical populations or older adults remains to be established. Additionally, while taxonomic shifts were identified, complementary functional analyses may be needed to fully elucidate the metabolic pathways involved and their relevance to host physiology. Furthermore, although diet logs were collected, the reliance on self-reported data posed a significant limitation to the accuracy of our dietary analysis. Finally, biological variability, including factors such as menstrual cycle phase - which was not controlled for despite its known influence on the gut microbiome - may also have contributed to inter-individual differences and obscured subtle effects.

## Conclusion

This study shows that resistance training can beneficially shape the gut microbiome, especially in individuals with marked strength gains. The enrichment of health-promoting taxa like *Faecalibacterium* and *Roseburia hominis* suggests potential anti-inflammatory and metabolic effects. However, these microbial shifts were not mirrored in the stool metabolome, indicating that functional outcomes may be subtle, localized, or require longer adaptation. Together with previous findings on endurance training, our results highlight physical activity—regardless of modality—as a promising, non-pharmacological approach to support metabolic and immune health. The consistent rise in SCFA-producing microbes across exercise types points to a shared mechanism, underscoring the value of multi-omics approaches in unraveling the complex interplay between exercise, the microbiome, and host physiology.

## Acknowledgments

NGS sequencing was performed with the support of the DFG-funded NGS Competence Center Tübingen (INST 37/1049-1) and the Institute for Medical Microbiology and Hygiene, University Hospital Tübingen. Data management and storage of raw data for this project were supported by the Quantitative Biology Center (QBiC), University of Tübingen, Germany. We are grateful to all study participants for their interest and effort as well as trainers for their enthusiasm to support and guide the study participants.

