## Supplemental Figures and Methods for "Resistance Training Reshapes the Gut Microbiome for Better Health"

**Supplementary Figures**

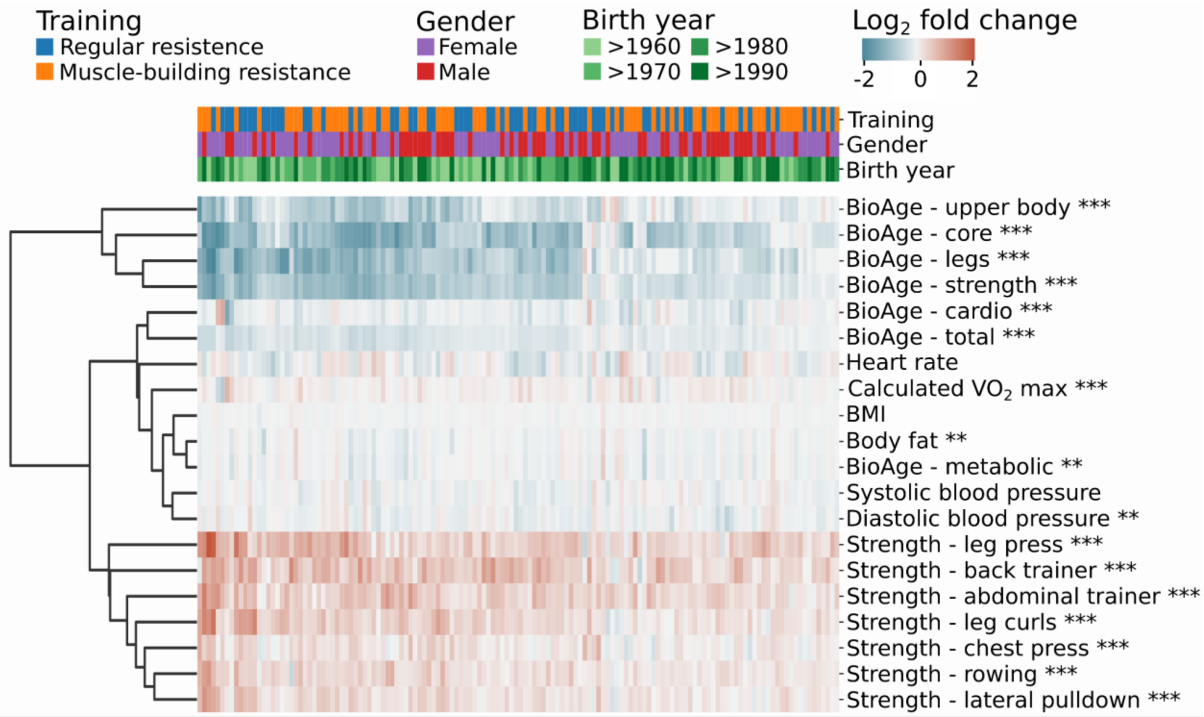


Suppl. Fig. 1: Heatmap of $\log_2$ fold changes from weeks 0 to 8 of fitness and health related metrics clustered hierarchically by euclidean distance with significance indicated by asterisks. Non-parametric Friedman test for repeated measures (formula: compound ~ weeks + (1|participant) ), Conover post hoc test, Benjamini-Hochberg adjusted p-values; *: p<0.05, **: p<0.01, ***: p<0.001

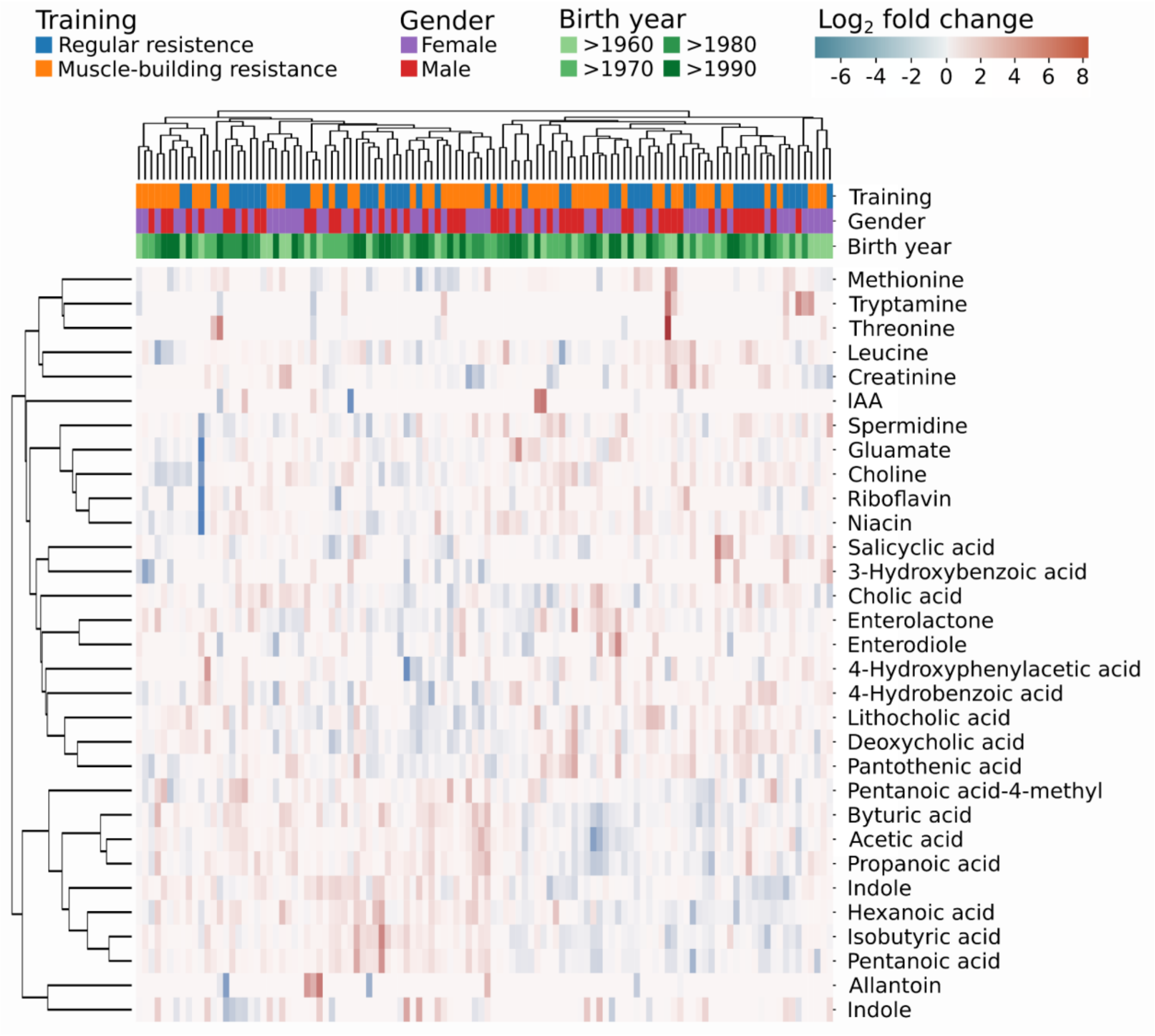


Suppl. Fig. 2: Heatmap of $\log_2$ fold changes from weeks 0 to 8 of metabolites clustered hierarchically by euclidean distance. No significant changes were detected using Friedman test adjusted with Benjamini-Hochberg to FDR or LME with model "metabolite ~ weeks + (1|Patient_id)".

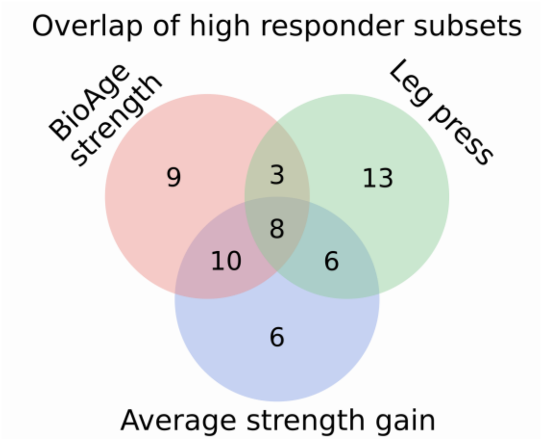


Suppl. Fig. 3: High responder (HR) subsets overlap partially. Participants were stratified into low-responder (LR, bottom 20%) and high-responder (HR, top 20%) across three strength metrics (Fig. 2C and Suppl. Fig. 4).

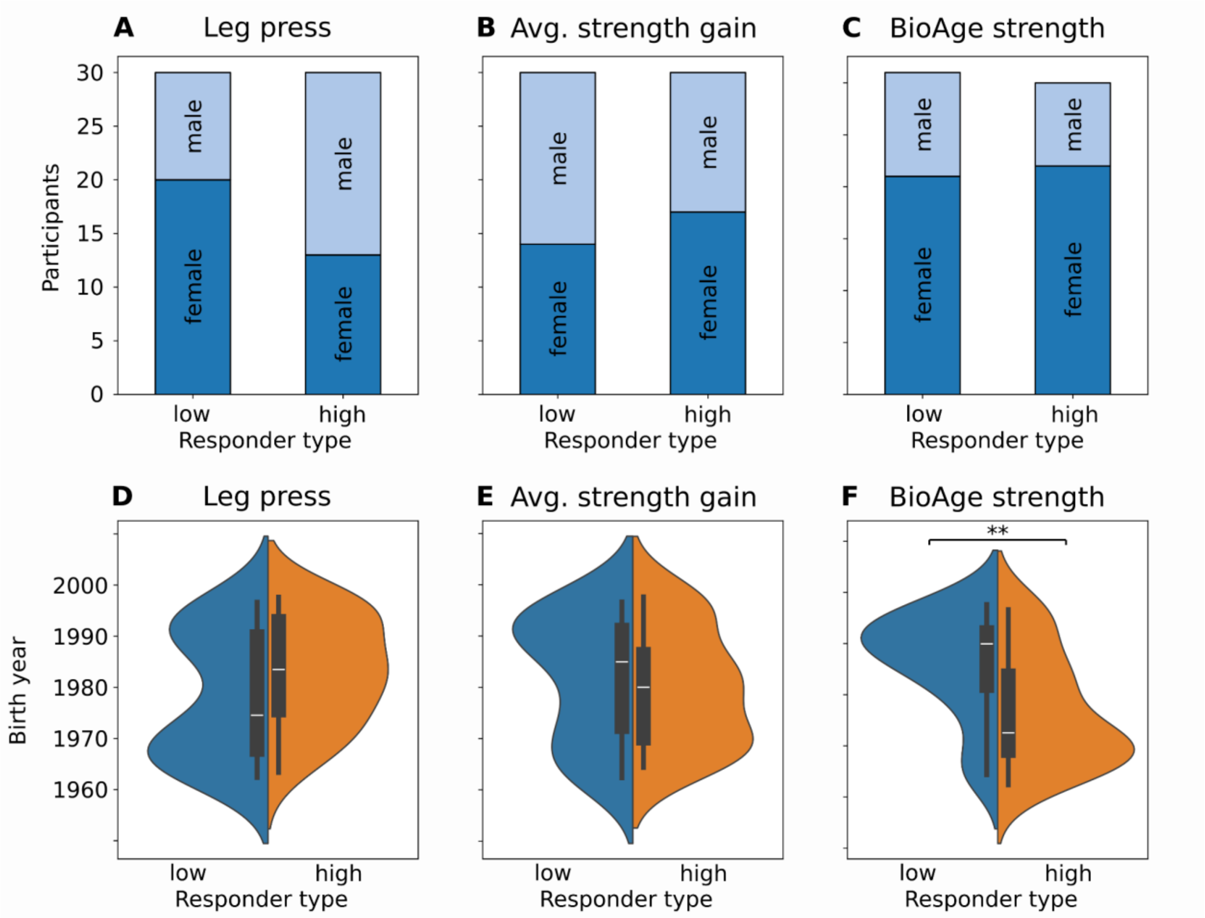


Suppl. Fig. 4: Demographics of low and high responders. Distribution of gender (A, B, C) and birth year (D, E, F) between low and high responder of maximum strength of leg press (A, D), average strength gain (B, E), or BioAge Strength (C, F). **: p≤0.01 using Kolmogorov-Smirnov test adjusted with Benjamini-Hochberg to FDR.

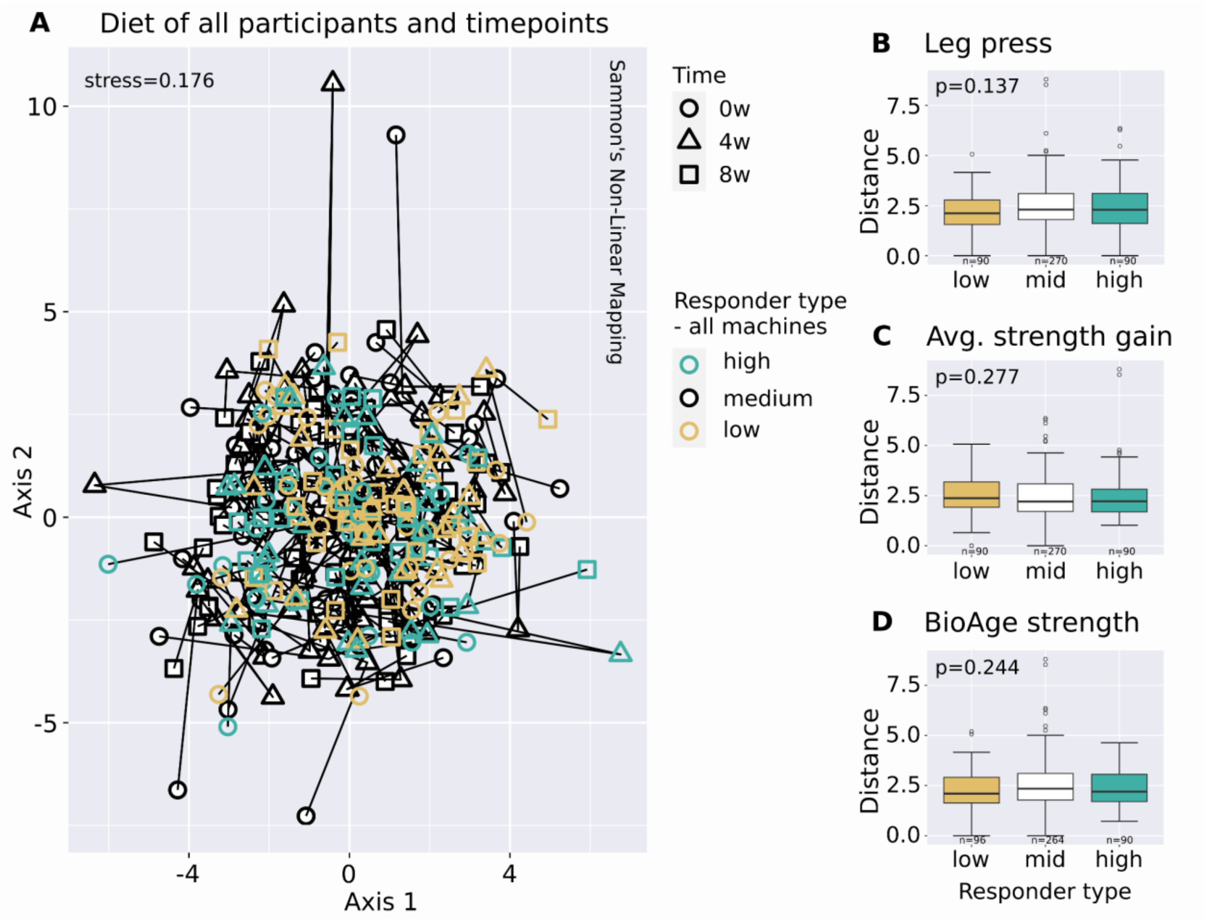


Suppl. Fig. 5: Diet differences within and between participants. (A) Ordination of summarized diet (food: fruit & vegetables, fish, meat, eggs, diary, grains, sweets, salty snacks; drinks: water & tea, coffee, juice & lemonade, alcohol in ml) based on euclidean distances of root-mean-square of centered values (R v4.2.3, package MASS v7.3-60). Lines connect timepoints per participant. Stress of <0.2 indicates a fair fit: usable representation, but higher values approach poor interpretation. (B, C, D) Diet distances within participants (0w-4w, 4w-8w, 0w-8w) stratified by responder type. p: Kruskal-Wallis rank sum test.

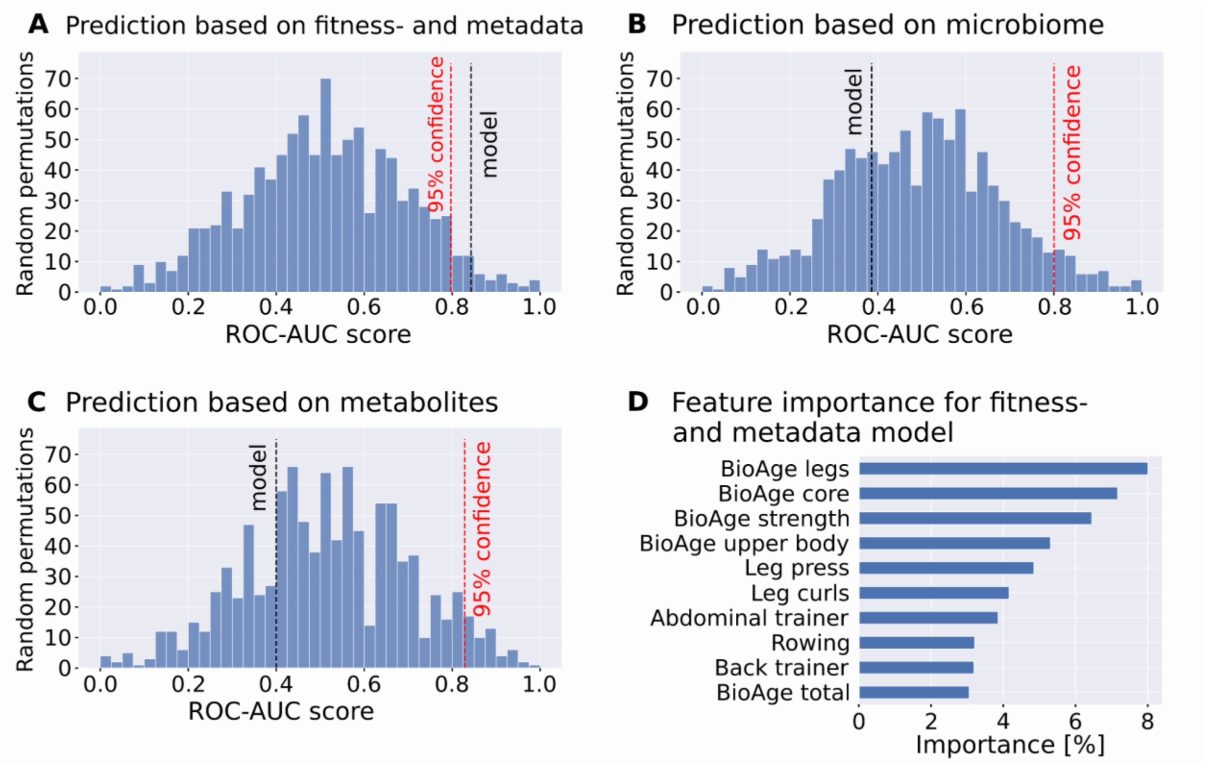


Suppl. Fig. 6: Prediction of low and high responder on all machines based on data collected at training start (baseline) with Random Forest. Model based on (A) fitness- and metadata - ROC-AUC=0.84, (B) microbiome - ROC-AUC=0.39, or (C) metabolites - ROC-AUC=0.40. (D) Ten most important features ("gini importance", https://doi.org/10.1201/9781315139470) in descending order of the prediction based on fitness- and metadata (i.e. model in A). Distribution of randomised training group predictions was achieved by permutation of the responder type (n=1000). Python v3.12.3 with scikit-learn v1.5.2.

**Supplementary Methods**

Strength and Training Metrics

The EGYM BioAge metric (referred to here as BioAge Total) is the average of three distinct BioAge components, including BioAge strength. In each of these, a lower BioAge indicates a healthier condition of the participant. Each BioAge component was calculated based on participant-specific data while accounting for sex differences. BioAge Metabolic is calculated based on body fat percentage, using a model informed by internal data and supported by previous literature describing the relationship between body fat and biological aging. BioAge Cardio is derived from estimated $VO_2$ max values, with sex-specific calculations. $VO_2$ max was estimated from maximal performance on cross trainers, using a model that combines internal performance data with previously validated methods [1–3]. The training instruments used are listed in Suppl. Table 1.

Suppl. Table 1: Instruments used

| Device | Measurements/Calculations | Company | Product |
|---|---|---|---|
| Blood pressure monitor | Resting diastolic & systolic blood pressure, pulse rate | A & D Medical | UA-767PC |
| Scale | Body weight, percent body fat, BMI | Seca | Seca Tru Fitness (seca 552) - 10000000671720 |
| Cross Trainer | $VO_2$ max | LifeFitness | Elevation Cross-Trainer - ASX127637 |
| SMart Strength machines | Maximum strength | EGYM | Smart Strength Generation 2 (2015) |

16S rRNA gene amplicon sequencing

NGS sequencing methods were performed at the Institute for Medical Microbiology and Hygiene (MGM) of the University of Tübingen. About 200mg of each stool sample were resuspended in 800 µL CD1 buffer (DNeasy 96 PowerSoil Pro QIAcube HT Kit, QIAGEN) and transferred to a ZR BashingBead Lysis Tube (Zymo Research). The tube was vortexed horizontally for 10 minutes on a vortex shaker. DNA was then purified using DNeasy 96 PowerSoil Pro QIAcube HT Kit (QIAGEN) according to the manufacturer's instructions.

Genomic DNA was quantified using Qubit dsDNA BR/HS Assay Kit (Thermo Fisher) and normalized to 30ng input for library preparation. The first step PCR was performed in 15µl reactions including KAPA HiFi HotStart ReadyMix (Roche), 515F and 806R primers (Caporaso et al. 2011) (~350 bp fragment of the 16S V4 region) and template DNA (PCR program: 95°C for 3 min, 28x (98°C for 20 sec, 55°C for 15 sec, 72° for 15 sec), 72°C for 5 min). First PCR products were purified using 12µl AMPure XP beads and eluted in 26µL 10mM Tris-HCl. Indexing was performed in the second step PCR including KAPA HiFi HotStart ReadyMix (Roche), index primer mix (IDT for Illumina DNA/RNA UD Indexes, Tagmentation), purified first PCR product as template (PCR program: 95°C for 3 min, 8x (95°C for 30 sec, 55°C for 30 sec, 72°C for 30 sec), 72°C for 5 min). After another bead purification (14µl AMPure XP beads, eluted in 15µL 10mM Tris-HCl) the libraries were checked for correct fragment length on agarose gels, quantified using QuantiFluor dsDNA System (Promega) and pooled equimolarly. The pools were sequenced on an Illumina MiSeq device with v2 sequencing kits with 2 x 250 bp read length and a depth of 2–156k reads per sample (8 samples <10k reads/sample, 9 samples >10k & <20k reads/sample, average 69k reads/sample).

16S rRNA gene amplicon analysis

Data processing, including quality control, reconstruction of sequences, taxonomic annotation, and diversity analysis was done using nf-core/ampliseq version 2.7.0 (doi: 10.5281/zenodo.10022435) [4] of the nf-core collection of workflows [5], utilising reproducible software environments from the Bioconda [6] and Biocontainers [7] projects.

The pipeline was executed with Nextflow v23.10.0 [8] and singularity v3.8.7 [9] with the following command: „`NXF_VER=23.10.0 nextflow run nf-core/ampliseq -r 2.7.0 -profile cfc --input samplesheet.tsv --FW_primer GTGYCAGCMGCCGCGGTAA --RV_primer GGACTACNVGGGTWTCTAAT --sample_inference pooled --trunclenf 200 --trunclenr 150 --min_len_asv 250 --max_len_asv 255 --cutadapt_min_overlap 15 --dada_ref_taxonomy silva=138 --skip_ancom --metadata metadata.tsv --outdir results`“. Data quality was evaluated with FastQC v0.12.1 [10] and summarized with MultiQC v1.15 [11]. Cutadapt v3.4 [12] trimmed primers and all untrimmed sequences were discarded, because sequences that did not contain primer sequences were considered artifacts. Less than 7.5% of the sequences were discarded per sample and a mean of 96.9% of the sequences per sample passed the filtering. Adapter and primer-free sequences were processed as one pool with DADA2 v1.28.0 [13] to eliminate PhiX contamination, trim reads (forward reads at 200 bp and reverse reads at 150 bp, reads shorter than this were discarded), discard reads with > 2 expected errors, correct errors, merge read pairs, and remove polymerase chain reaction (PCR) chimeras. Ultimately,

9057 amplicon sequencing variants (ASVs) were obtained across all samples. The ASV count table contained in total 27,253,674 counts, at least 2,043 and at most 140,951 per sample (average 60,564). To remove spurious sequences, 7 ASVs with length lower than 250 or above 255 bp were removed with less than 0.42% counts per sample (9050 ASVs passed). Taxonomic classification was performed by DADA2 and the database 'Silva 138.1 prokaryotic SSU' [14]. ASV sequences, abundance and DADA2 taxonomic assignments were loaded into QIIME2 v2023.7.0 [15]. 19 ASVs that were annotated as mitochondria were discarded (9,031 ASVs passed). Within QIIME2, the final microbial community data was investigated for alpha (within-sample) and beta (between-sample) diversity after rarefaction to 2,042 counts.

Metabolome analysis

Metabolomics sample preparation and measurements were performed at The Center for Plant Molecular Biology (ZMBP) of the University of Tübingen. Frozen samples were thawed and fully transferred to 5 mL reaction tubes. Tara weights and sample weights were documented prior to subsequent processing. Samples were then centrifuged using a Hettich swing out centrifuge for 10 minutes at 4500 rpm. The supernatant was transferred to new 2 mL reaction tubes and 1 ml was prepared for LC-MS, while the remaining volume was used for GC-MS. Resulting pellets were left for 20 minutes in the fume hood, then frozen at -80°C before freeze drying for 48 hours.

The LC-MS aliquot was dried using a speed vac, and pellet was resuspended using 200µL $H_2O$, 20% Methanol, 0.1% formic acid, with 9µM Lenk as internal standard. Samples were subjected to ultrasound for 10 minutes, 10 minutes incubation at room temperature, vortexing and again 10 minutes incubation at room temperature. Subsequently, samples were centrifuged at 14000 rpm for 15 minutes at 4°C, and 100µL supernatants were transferred to autosampler and 3 µL were injected into LC-MS.
LC-MS analyses were carried out with a "Waters Acquity-SynaptG2" LC-MS system, operated at MS/MSE mode in parallel with a resolution of 10000, a scan time of 0.2 seconds and a scan range from 50 - 2000 m/z. For analyte separation an "Acquity UPLC HSS C18 SB, 2.1x100mm, 1.8µm" column was used with a 10 min gradient from Water with 0.1% Formic Acid (Solution A) to Methanol with 0.1% Formic Acid (Solution B).
GC-MS aliquots were centrifuged at 13000 rpm for 10 minutes at 4°C. Supernatant was transferred to 10 mL Headspace screwcap vials according to pipetting scheme Suppl. Table 2. Samples were measured on Shimadzu TQ8040 equipped with headspace sampler HS20, running in Headspace LOOP mode, settings are detailed in Supplemental File 1). Column Stabilwax-DA (Restek), length 30 m, diameter 0.32 mm, film thickness 1.0 µm was used. Peaks were checked manually and peak integration was performed using Labsolutions Insight

GC/MS, while relative peak areas and metabolite amounts were calculated using R. Data was normalized using internal standards and stool concentration, calculated by the dry weight of stool in the total amount of stool sample given and all subsequent analyses were performed on normalized quantities of the metabolites.

Suppl. Table 2: Pipetting scheme for GC-MS samples. ISTD: Internal standard

| Component | Volume [µL] |
|---|---|
| Saltout (contains 882 $gL^{-1}$ of $(NH_4)_2SO_4$, and 238 $gL^{-1}$ of $NaH_2PO_4$ | 3800 |
| ISTD: 200 µM Butyric acid d8 (equals 20 nmol/Sample) | 100 |
| Sample | 150 |
| $H_3PO_4$ | 450 |
| TOTAL | 4500 |
